# Progeny-based genomic selection reveals untapped genetic potential in an underutilized medicinal plant, *Perilla frutescens*

**DOI:** 10.1101/2025.11.26.690889

**Authors:** Sei Kinoshita, Kengo Sakurai, Takahiro Tsusaka, Miki Sakurai, Kenta Shirasawa, Sachiko Isobe, Hiroyoshi Iwata

**Affiliations:** Graduate School of Agricultural and Life Sciences, University of Tokyo, Tokyo, Japan; TSUMURA & CO., Ibaraki, Japan; LAO TSUMURA CO., LTD., Salavan, Laos; Kazusa DNA Research Institute, Chiba, Japan

**Keywords:** Crossing experiment, genome-assisted breeding, genomic selection, medicinal plants, *Perilla frutescens*

## Abstract

- Despite their substantial therapeutic value, medicinal plants have undergone limited genetic improvement through breeding because of the scarcity of expert breeders. Moreover, quantifying bioactive compounds is expensive. Genomic selection (GS), which leverages genome-wide markers to predict breeding values and assemble favorable alleles, offers a practical way to unlock latent genetic potential. As a model case, we evaluated GS in red perilla (*Perilla frutescens*).
- Building on previous work, we implemented a cross-selection strategy that prioritized segregation variance by selecting crosses based on predicted additive genotypic values of the progeny, and evaluated its effectiveness through actual crossing experiments targeting three key medicinal compounds.
- Progeny from GS-based crosses (Crs1–Crs7) outperformed those from phenotypic selection (Crs8) in the G_2_ generation, demonstrating a higher mean, greater variance, and superior top individuals. The best G_2_ individual exhibited nearly twofold higher levels of two target compounds relative to the existing cultivar ‘Sekiho’.
- This study provides the first empirical demonstration that GS can improve multiple medicinal compounds in red perilla and highlights the effectiveness of cross-selection based on predicted progeny performance. In addition, the evidence presented here supports the broader application of GS in underutilized medicinal plants.

## Introduction

Bioactive compounds naturally derived from plants have been used to restore and support human health since ancient times. Recently, medicinal and herbal plants have gained renewed global attention. According to the World Health Organization (WHO), approximately 80% of the world’s population uses plant-based medicines, and traditional plant-derived remedies are employed in 170 WHO member countries (Zamani et al., 2025; World Health Organization (WHO), 2025). Today, medicinal and herbal plants are not only used for treating diseases, but are also increasingly incorporated into dietary supplements for health maintenance and enhancement. Furthermore, reflecting the growing consumer preference for products that are perceived as natural rather than synthetic, these plants are widely used in natural cosmetics and personal care. Reflecting this growing demand, the global market for medicinal and herbal plants reached USD 215.44 billion in 2024 and is projected to expand to USD 403.50 billion by 2033 (Zamani et al., 2025; Market Data Forecast, 2025). The development of superior cultivars is essential to ensure a stable supply of high-quality raw plant materials (Hansan et al., 2024). Despite their increasing demand and economic importance, research on medicinal and herbal plants has predominantly focused on identifying and characterizing their bioactive compounds, whereas breeding-oriented studies remain scarce.

Two major challenges exist in breeding medicinal and herbal plants. First, these plants are generally considered underutilized crops, and, unlike major crops, a few experienced breeders are capable of selecting superior individuals based solely on their phenotypic traits. Secondly, target traits and bioactive medicinal compounds are difficult to evaluate visually, and their quantification for selection purposes is both time-consuming and costly (Hansan et al., 2024; Akpojotor et al, 2024). In contrast to major crops, for which breeding has already achieved substantial progress, medicinal and herbal plants remain largely unimproved. Therefore, these species have great potential for genetic improvement, and genome-based breeding approaches offer promising opportunities to accelerate the development of superior cultivars (Kusmec et al., 2021). To address these challenges, our research group considered genomic selection (GS, Meuwissen et al., 2001), a selection method that estimates the genomic breeding values of individuals based on genome-wide marker information, as a promising approach for breeding medicinal plants (Meuwissen et al., 2001; Kinoshita et al., 2023). Since its proposal by Meuwissen et al. (2001), GS has been successfully applied to a wide range of plant species. Its effectiveness has been demonstrated not only in major crops, such as rice, maize, and wheat, but also in underutilized crops, such as buckwheat. In addition, experimental studies have demonstrated that GS can outperform conventional phenotypic selection (Beyene et al., 2015; Yabe et al., 2018; Beyene et al., 2019; Xu et al., 2021). To the best of our knowledge, no empirical studies have demonstrated the effectiveness of GS in medicinal plants.

We used *Perilla frutescens* (L.) (“shiso” in Japanese), a self-pollinating annual plant belonging to the Lamiaceae family, to establish a model case for GS-based breeding of medicinal plants. We selected *Perilla frutescens* because it has a short generation cycle, a relatively small genome (∼1.25 Gb; Tamura et al., 2023) that facilitates whole-genome sequencing and genetic analysis, and simple reproductive biology as a self-pollinating species. Notably, *Perilla frutescens* is also in substantial demand. It has been traditionally cultivated and utilized in East Asian countries, particularly China, Japan, and Korea (Nitta, Lee & Ohnishi, 2003; Dhyani, Chopra & Garg, 2019). *Perilla frutescens* has diverse applications, and its leaves, stems, and seeds are used for medicinal, culinary, and cosmetic purposes (Wu et al., 2023). As a medicinal plant, *Perilla frutescens* is used as a raw material in traditional medicine, including Kampo in Japan. *Perilla frutescens* contains several beneficial bioactive constituents, including volatile oils, flavonoids, and phenolic acids. It is used in Kampo medicine as an ingredient in traditional formulations, such as Hangekobokuto and Kososan, which are effective for a range of symptoms, including anxiety, cold, and gastrointestinal weakness (Igarashi & Miyazaki, 2013; Yi, Wang & Peng, 2025). Among the several bioactive constituents of *Perilla frutescens*, perillaldehyde, rosmarinic acid, and anthocyanins are considered key components of Kampo medicine (Ogawa, Hikosaka & Goto, 2018). Perillaldehyde is an essential oil that has been reported to possess sedative antifungal, antibacterial, and antioxidant properties (Honda et al., 1986; Yu et al., 2017; Erhunmwunsee et al., 2022). Rosmarinic acid, a type of phenolic compound, is known for its antidepressive, anti-inflammatory, antioxidant, and antitumor properties and has been reported to be effective against conditions such as Alzheimer’s disease and allergies (Takeda et al., 2002; Swamy, Sinniah & Ghasemzadeh, 2018; Adam et al., 2023). Moreover, anthocyanin is a type of polyphenol with pharmacological effects such as antioxidant activity. In addition to its bioactivity, it serves as a pigment responsible for the purple coloration of *Perilla* leaves and contributes to the appearance of Kampo formulations (Asif, 2012; Hou et al., 2022).

Our breeding goal is to develop superior red perilla lines with higher levels of the three major medicinal compounds—perillaldehyde, rosmarinic acid, and anthocyanin—compared to the existing cultivar ‘Sekiho’. To support this objective, we first evaluated the genetic characteristics of major medicinal compounds in Perilla and the effectiveness of GS by conducting QTL analysis and genomic prediction (GP) (Kinoshita et al., 2023). All three traits—perillaldehyde, rosmarinic acid, and anthocyanin—exhibited moderate-to-high heritability. In addition, the prediction accuracy of GP ranged from 0.48 to 0.85, indicating that reasonably high accuracy was achieved across traits. Moreover, three biparental populations exhibited superior traits (Kinoshita et al., 2023). This indicates that continuing selection within each population may only improve a single trait. Therefore, we hypothesized that crossing populations, while accounting for their distinct genetic effects, could facilitate the accumulation of favorable alleles from each population. We tested this hypothesis using simulation studies (Kinoshita et al., 2025). We adopted a strategy that emphasizes genetic variance to select cross-pairs (Bernardo, 2014). The additive genotypic values of the progeny were predicted via simulations using this approach, allowing us to identify crosses capable of producing superior progenies. The development of new cultivars for plant breeding is achieved through iterative cycles of selecting superior individuals and crossing. Thus, identifying superior crossing pairs is critical. This is particularly true for underutilized medicinal plants, where a few elite lines have accumulated multiple favorable alleles and genetic improvement necessarily depends on the assembly of useful alleles through crossing. The selection strategy we employed, explicitly accounting for the genetic variance of progeny, helps maintain genetic diversity and has been demonstrated by numerous simulation studies, as well as several empirical studies on major crops, to promote medium-to long-term genetic gain (Iwata et al., 2013; Bernardo, 2014; Rembe et al., 2022; Miller et al., 2023). Nevertheless, empirical validations of cross-selection based on the predicted genetic ability of progeny remain scarce, and to the best of our knowledge, no such demonstration exists for medicinal plants.

In this study, we evaluated the effectiveness of GS in Perilla, particularly the utility of cross-selection based on the predicted additive genotypic values of the progeny. We compared this GS-based approach with conventional phenotypic selection by conducting actual crossing experiments and assessed the extent to which the medicinal compound content was improved relative to that of the existing red perilla cultivar, ‘Sekiho’.

## Materials and Methods

### Plant materials for breeding experiment

A schematic representation of the breeding experiment is shown in Fig. 1. The initial breeding materials comprised three biparental populations, each of which advanced to the F_5_ generation through selection and selfing. Subsequently, inter- and intrapopulation crosses among these three populations were performed to generate progenies with mixed genetic backgrounds. The original biparental populations were derived from the following crosses: ‘Sekiho’ × ‘st27’, ‘Sekiho’ × ‘st40’, and ‘Sekiho’ × ‘st44’. Hereafter, these are referred to as S827, S840, and S844, respectively. ‘Sekiho’ is a red perilla variety owned by TSUMURA & CO., whereas ‘st27’, ‘st40’, and ‘st44’ are green perilla genetic resources maintained by the National Agriculture and Food Research Organization (NARO). Taxonomically, ‘Sekiho’, ‘st27’, and ‘st44’ are classified as *Perilla frutescens* (L.) Britton var. *crispa*, whereas ‘st40’ is classified as *P. frutescens* (L.) Britton var. *frutescens*.

**Figure 1.**
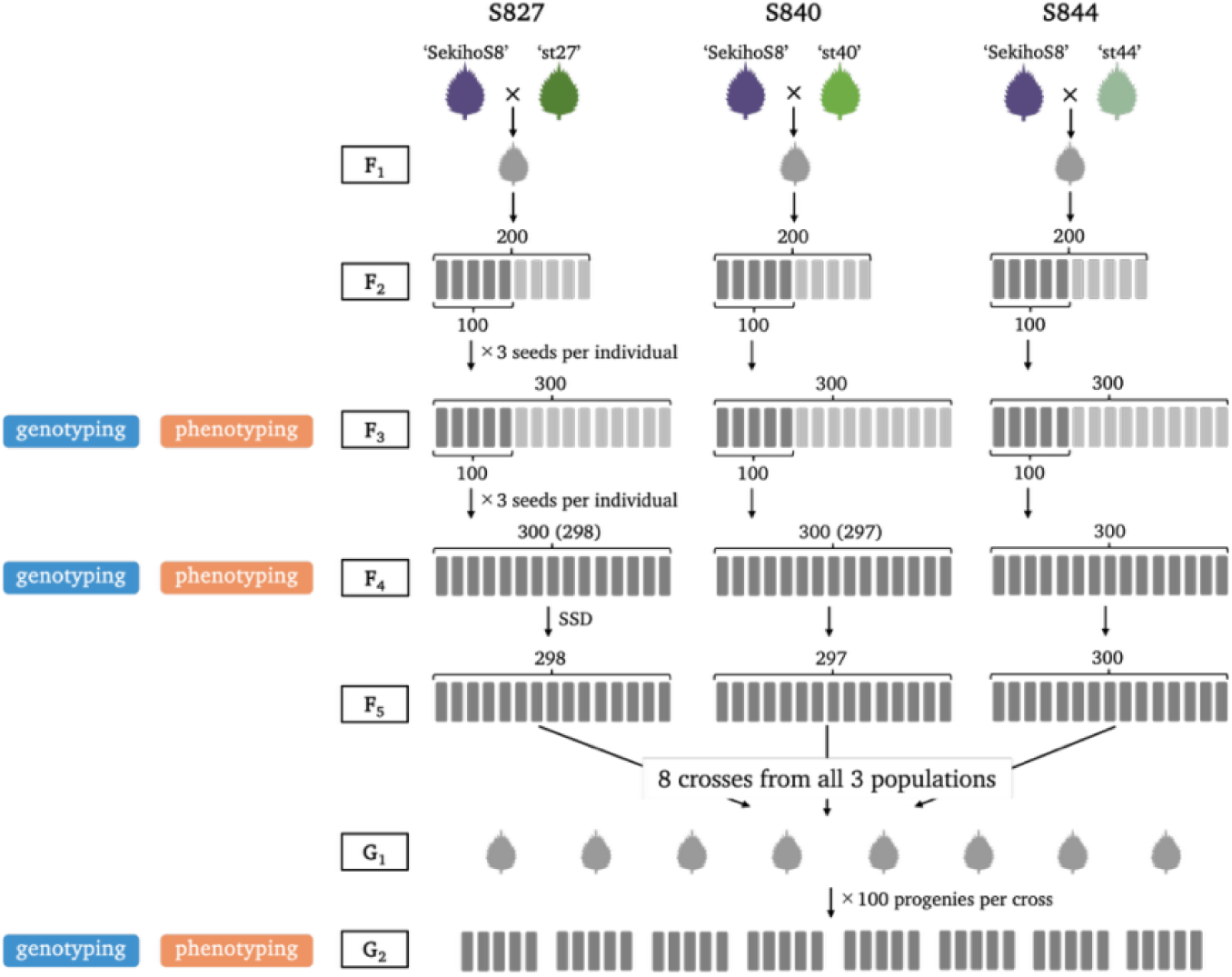
A schematic representation of the breeding experiment. Three biparental populations (S827, S840, and S844) were developed and advanced to the F₅ generation through selection and selfing. Inter- or intra-population crosses were subsequently conducted to produce the selfed second generation after crossing (G₂). Genotypic and phenotypic data were collected from the F₃, F₄, and G₂ generations.

Inter- or intra-population crosses were subsequently conducted to produce the selfed second generation after crossing (G₂). Genotypic and phenotypic data were collected from the F₃, F₄, and G₂ generations.

From each of the three biparental populations, 200 F_2_ individuals were grown in the field. Of these, 100 individuals per population were randomly selected, and three seeds from each plant were sown to produce the F_3_ generation. From the 300 F_3_ plants per population grown in the field, 50 individuals were randomly selected. Another 50 were selected from the F_3_ individuals that had not been cultivated in the field, and three seeds per plant were used to produce the F_4_ generation. Seeds from uncultivated F_3_ individuals were included to preserve genetic diversity. From the F_4_ to F_5_ generations, all the individuals were advanced using the single-seed descent (SSD) method. A total of 895 F_5_ individuals were obtained from the three populations (298 from S827, 297 from S840, and 300 from S844). From these, eight pairs (comprising seven individuals) were selected and crossed to generate the G_1_ generation. The criteria and method for selecting these eight crosses are described in the section “*Selection of cross pairs in the F_5_ generation*.” For the G_2_ generation, 100 individuals from each of the eight crosses were grown in the field, resulting in 800 G_2_ individuals evaluated under field conditions.

### Genotype and phenotype data

Marker genotype and phenotype data were collected from the F_3_, F_4_, and G_2_ generations. The number of individuals for which both genotype and phenotype data were obtained was as follows: 900 individuals in the F_3_ generation (300 from each population), 895 individuals in the F_4_ generation (298 from S827, 297 from S840, and 300 from S844), and 800 individuals in the G_2_ generation. In addition, ‘Sekiho’ was grown alongside the F_3_, F_4_, and G_2_ generations in the same field, and phenotypic data were collected for each generation.

Procedures for genomic DNA extraction and variant filtering were consistent across all generations. Genomic DNA was extracted using the double-digestion restriction site-associated DNA sequencing (dd-RAD-seq) method (Shirasawa, Hirakawa & Isobe, 2016). The obtained reads were mapped to the assembled sequence of ‘Sekiho’ using Bowtie2 (Langmead & Salzberg, 2012; Kinoshita et al., 2025). Variant calling and quality control were conducted with VCFtools version 0.1.16, using a max-missing threshold of 0.9 (Danecek et al., 2011). Single-nucleotide polymorphism (SNP) markers with a minor allele frequency (MAF) below 0.025 were excluded, and missing genotypes were imputed using Beagle 5.1 (Browning, Zhou & Browning, 2018). Subsequently, MAF was independently applied to each generation. In the F_3_ and F_4_ generations, SNPs were filtered within each population and across all three populations using a threshold of MAF < 0.01. In the G_2_ generation, the SNPs were filtered across the entire dataset using the same threshold. Thus, 1,964 SNPs remained in the F_3_ generation when all individuals were considered, with 727, 1,432, and 579 SNPs retained in the S827, S840, and S844, respectively. The F_4_ generation had the same total number of SNPs (1,964), with 862, 1,432, and 579 SNPs at S872, S840, and S844, respectively. In the G_2_ generation, 1,297 SNPs remained after filtering. For subsequent analyses, genotype scores were encoded as follows: homozygous SNPs identical to ‘Sekiho’ were assigned a score of 0, heterozygous SNPs a score of 1, and homozygous SNPs differing from ‘Sekiho’ a score of 2.

Among the target traits, perillaldehyde, rosmarinic acid, anthocyanin—perillaldehyde and rosmarinic acid contents were measured in the F_3_, F_4_, and G_2_ generations, whereas anthocyanin content was measured only in the F_3_ and G_2_ generations. The procedures for phenotypic data collection in the F_3_ and F_4_ generations, as well as for the quantification of perillaldehyde and rosmarinic acid contents in the G_2_ generation, followed the methods described by Kinoshita et al. (2023). Because the procedure for measuring the anthocyanin content in the G_2_ generation differed slightly from that used in the F_3_ and F_4_ generations, the specific methodology is described below. For anthocyanin quantification, 0.2 g of powdered leaf sample was placed in 30 mL of 3% trifluoroacetic acid (TFA), shaken at 200 rpm for 10 min, and centrifuged at 3,000 rpm for 10 min. The supernatant was collected, and the residue was re-extracted with 15 mL of 3% TFA using the same procedure. Supernatants were combined and diluted to 50 mL with 3% TFA. The extract was filtered through a 0.45 µm membrane filter. For measurement, 500 µL of the filtrate was mixed with 1 mL of 3% TFA, and the absorbance was measured at 520 nm using a NanoDrop spectrophotometer in UV–Vis mode with a 10 mm cuvette pathlength.

### Genomic prediction model

As described in Kinoshita et al. (2023), Genomic prediction models (GBLUP and BayesB) were constructed for each of the three medicinal compounds in each population of the F_3_ and F_4_ generations, as described by Kinoshita et al. (2023). These models were used to evaluate the genomic prediction accuracy and estimate genomic heritability. According to the results reported by Kinoshita et al. (2023), the BayesB model achieved comparable or higher prediction accuracy than the GBLUP model. Therefore, marker effects estimated using the BayesB model were used for selection. A reference genome assembly different from that used by Kinoshita et al. (2023) was used for genotyping. The genomic prediction accuracy based on the Bayes B model and genomic heritability estimated by the GBLUP model are provided in the Supplementary File (Fig. S1). The observed trends were consistent with those previously reported by Kinoshita et al. (2023). The GBLUP model was implemented using the R package “rrBLUP” (version 4.6.3; Endelman, 2011), and the BayesB model was implemented using the R package “BGLR” (version 1.1.3; Pérez & De Los Campos, 2014).

### Selection of cross pairs in the F_5_ generation

In the F_5_ generation, seven cross pairs (Crs1–Crs7) were selected based on the additive genotypic values of their simulated progenies (G_2_ generation). In addition, one cross (Crs8) was selected based on phenotypic values observed in the F_4_ generation. Among individuals that exhibited higher levels of both perillaldehyde and rosmarinic acid than ‘Sekiho’, the individual with the highest perillaldehyde content and the one with the highest rosmarinic acid content were crossed in the F_5_ generation to form Crs8.

The procedure for selecting Crs1–Crs7 is described in detail below. Using marker genotype data from the F_4_ generation and marker effects estimated separately for each population, simulations were conducted according to the scheme shown in Fig. 2. In the simulation, individuals were advanced from the F_4_ to F_5_ generations through the SSD. In the F_5_ generation, selected individuals were crossed to produce four G_1_ individuals per cross. Next, each G_1_ individual was selfed to produce 500 G_2_ individuals, resulting in 2,000 simulated G_2_ individuals per cross. Additive genotypic values were calculated for all the simulated G_2_ individuals within each cross. The top 1% of individuals (i.e., the 20th-ranked individual) based on the additive genotypic value was used as the performance metric for a single cross pair and a single simulation run. This process was repeated 500 times, and the average of the 500 values was used as the predicted performance score for each cross.

**Figure 2.**
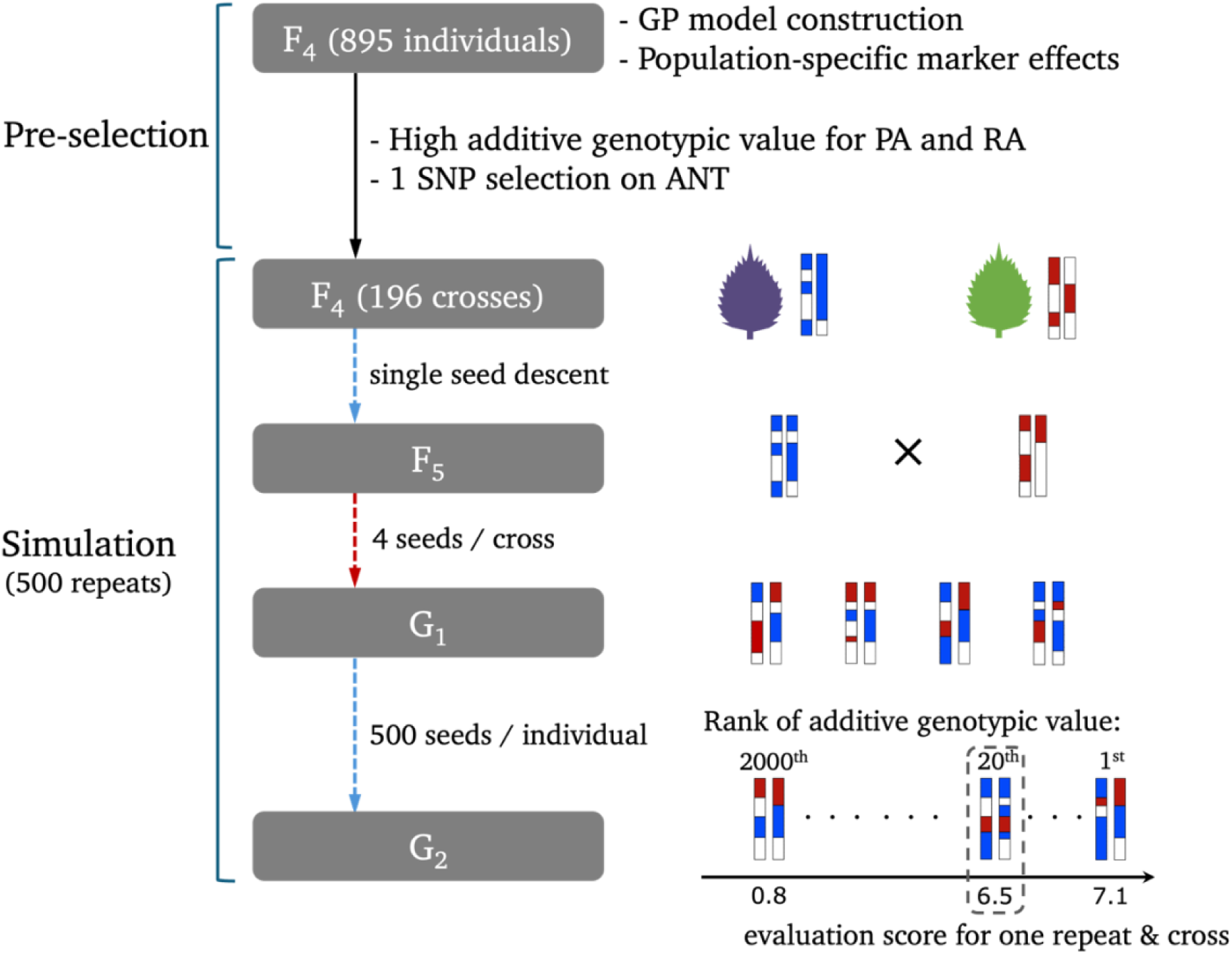
Workflow of the simulation for cross selection based on predicted progeny performance. To reduce computation time, pre-selection was performed on the F₄ generation, and simulations were conducted for 196 selected cross pairs using the F₄ genotypic data. The schematic on the right illustrates the conceptual flow of the simulation process for one cross and one replication.

To reduce computational time, the simulation described above was not performed for all possible cross pairs among the 895 F_4_ individuals. Instead, evaluation values were calculated for a subset of 196 pairs selected using the procedure described below. First, the S844 population was excluded from the analysis because it contained a smaller number of SNPs and exhibited lower genomic prediction accuracy than the other two populations. Consequently, cross pairs were evaluated among 595 F_4_ individuals from the S827 and S840 populations. Among the 176,715 possible pairs derived from these two populations, the selection index 𝑢̅, was computed for each cross. The index value 𝑢̅_𝑘_ for cross *k,* composed of individuals i and j, is defined as follows:

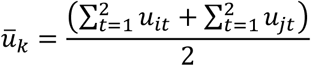

where 𝑢_𝑖𝑡_ and 𝑢_𝑗𝑡_ represent the additive genotypic values of individuals *i* and *j*, respectively, for the t^th^ trait (t = 1 for perillaldehyde; t = 2 for rosmarinic acid). These values were calculated using the marker effects estimated within the population to which each individual belonged. In addition, 𝑢_𝑖𝑡_ and 𝑢_𝑗𝑡_ were scaled on a per-trait basis before their use in the index calculations. Based on the calculated 𝑢̅_𝑘_ for all 176,715 possible cross pairs, the top 1,000 pairs with the highest index values were selected. To ensure anthocyanin expression in the progeny, these 1,000 pairs were further filtered using a marker on chromosome 8, which was previously identified by Kinoshita et al. (2023) as associated with anthocyanin accumulation. Specifically, only cross pairs in which both parental individuals carried the ‘Sekiho’ allele (genotype score = 0) at this marker were retained.

Consequently, 196 cross pairs remained, and the simulation procedure described above was applied to the selected pairs.

The predicted additive genotypic values in the G_2_ generation based on these simulations are shown in Fig. 3. Among the 196 selected cross pairs, 116 were intra-population crosses from the S827 population, 5 were intra-population crosses from the S840 population, and 75 were inter-population crosses between S827 and S840. For each of the three cross types, Pareto frontiers were constructed based on the predicted additive genotypic values of the G_2_ generation. The construction of Pareto frontiers was performed using the R package “rPref” (version 1.4.0). In Fig. 3, the colored dots represent the predicted values of the G_2_ individuals derived from the cross-pairs identified as Pareto optimal. Based on the Pareto frontier, two intra-population crosses from S827 (Crs1 and Crs7), one intra-population cross from S840 (Crs6), and four inter-population crosses (Crs2–Crs5) were selected — chosen in consultation with the breeder — for crossing to produce the G_2_ generation. The individuals used to construct the eight crosses are listed in Table S1. Crs1–Crs7 comprised five individuals: three from the S827 population and two from the S840 population. In contrast, Crs8 was composed of two different individuals, one from the S827 population and the other from the S844 population.

**Figure 3.**
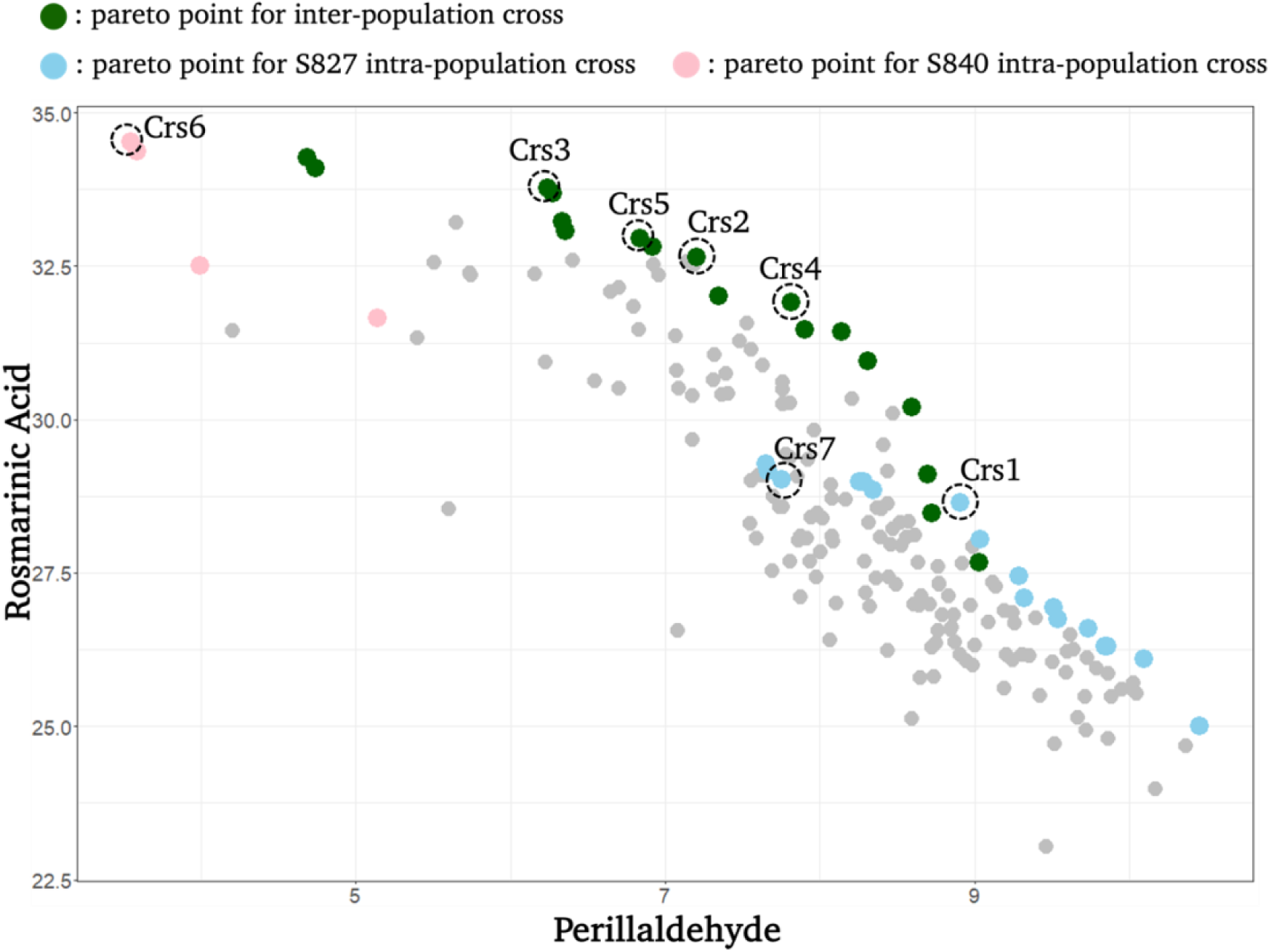
Prediction of additive genotypic value in G₂ progeny based on marker genotype data of the F₄ generation obtained by simulation. Each point represents the predicted additive genotypic value of the progeny derived from one of the 196 F₄ crosses. Dark green, sky blue, and pink points indicate the pareto-optimal crosses derived from inter-population crosses, intra-population crosses within S827, and intra-population crosses within S840, respectively. Points enclosed by dashed circles correspond to the crosses actually used to produce the G₂ generation. Labels Crs1–7 denote the names of the specific crosses used in this study.

### Inference of allele origin in the G_2_ generation

Marker effects were estimated separately for the S827 and S840 populations and used to select G_2_ parental individuals under the GS framework. This approach allowed the GS-based selection to account for the effects of three allelic types—those derived from ‘Sekiho’, ‘st27’, and ‘st40’. Accordingly, the origin of alleles (i.e., ‘Sekiho’, ‘st27’, or ‘st40’) in the G_2_ generation was inferred for each individual. The method used to infer the allele origin is illustrated in Fig. S2.

For each individual in the F_4_ generation, haplotypes were coded as 0 or 1: 0 representing alleles derived from ‘Sekiho’, and 1 representing alleles derived from either ‘st27’ or ‘st40’, depending on the population to which the individual belonged. Crossing was performed in the F_5_ generation after one generation of selfing from F_4_, and allele origins were inferred from the resulting G_2_ progeny. Here, alleles derived from ‘Sekiho’, ‘st27’, and ‘st40’ are denoted as A, B_1_, and B_2_, respectively. For example, if the genotypes at a given SNP in two F_4_ parents are (AA, B_1_B_1_), this indicates that one individual is homozygous for ‘Sekiho’ and the other is homozygous for ‘st27’. In the G_2_ generation, the allele origin can be clearly inferred under the following conditions:

1. The genotype at a given SNP in one G_2_ individual is homozygous for ‘Sekiho’ or
2. Both F_4_ parents carry the same B-type allele at that SNP.

These conditions included the following genotype combinations in the F_4_ parents: (AA, AB_1_), (AB_1_, AB_1_), (AA, B_1_B_1_), (AB_1_, B_1_B_1_), and (B_1_B_1_, B_1_B_1_), with equivalent logic applied when B_1_ was replaced with B_2_. Additionally, when the F_4_ parents carry different B-type alleles, as in the case of (AB_1_, B_2_B_2_), the origin of the G_2_ genotype can still be determined if the G_2_ genotype is either homozygous for ‘Sekiho’ or heterozygous. In contrast, if the F_4_ genotype is (AB_1_, AB_2_) or (B_1_B_1_, B_2_B_2_), the allele origin of the G_2_ genotype cannot be determined with certainty (Fig. S2). Although hidden Markov models or flanking markers can be used to probabilistically infer allele origins in such ambiguous cases, the proportion of these markers was small in this study. Therefore, we assigned allele origins only to markers whose origins could be determined with certainty using the above rules and treated all other cases as missing.

Using the markers for which allele origins were determined using the method described above, together with the marker effects estimated separately for each population in the F_4_ generation using the Bayes B model, we predicted the additive genotypic values of G_2_ individuals for each trait using the following equation:

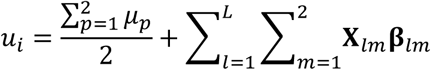

where 𝑢_𝑖_ represents the predicted additive genotypic value of the 𝑖^th^ individual in the G_2_ generation. 𝜇_𝑝_ denotes the population mean estimated from the F_4_ generation, corresponding to the population(s) from which the two parents originated. In the case of inter-population crosses (e.g., Crs2–Crs5 and Crs8), the two parents originate from different populations, and thus, 𝜇_1_ ≠ 𝜇_2_; 𝜇_1_ and 𝜇_2_ are the population means estimated from the respective parental populations. 𝐗_𝑙𝑚_ represents the marker genotype matrix constructed based on the inferred allele origin, with dimensions 𝐿 × 2 or 1 (where 𝐿 = 1,297 is the number of markers, and the number of columns corresponds to the number of allele origins: two for inter-population crosses and one for intra-population crosses). Genotypes were coded as 0, 1, or 2. 𝛃_𝑙𝑚_ is a matrix of marker effects with the same dimensions as 𝐗_𝑙𝑚_, where each column corresponds to the marker effects estimated from one of the parental populations. For markers with undetermined allele origins, the corresponding genotype scores were set to 0 so that they did not contribute to the prediction of additive genotypic values.

We next evaluated the effectiveness of incorporating population-specific marker effects, for which we compared the G_2_ additive genotypic values predicted using the parental model described above with those predicted using an alternative approach that did not distinguish the allele origins. In the latter approach, hereafter referred to as the bi-allelic model, only two allele types were considered for each SNP, corresponding to whether the allele matched or differed from the reference genome used for genotyping. Marker effects for the biallelic model were estimated using the BayesB model across all three populations combined in the F_4_ generation. In this study, we refer to the model that distinguishes allele origins and uses population-specific marker effects as the parental model and the model that does not distinguish allele origins as the bi-allelic model. The prediction accuracy of the two models for G_2_ generation was assessed using Pearson’s correlation between the predicted and observed values.

## Results

### Selected cross pairs in the F_5_ generation

Fig. 4 shows the perillaldehyde and rosmarinic acid phenotypes in the F_4_ generation of the seven individuals that constituted the eight selected crosses. Although the actual crosses were performed using the F_5_ generation, obtained by selfing these F_4_ individuals via SSD, phenotypic data were not collected in the F_5_ generation; therefore, data from the F_4_ generation are presented here. The five individuals selected based on the predicted additive genotypic values of the progeny (Crs1 to Crs7) are indicated by black circles, whereas the two individuals selected based on the phenotypic values in the F_4_ generation (Crs8) are indicated by black triangles. The two red lines represent the phenotypic values of ‘Sekiho’ grown in the same year. Among the five individuals selected for Crs1–Crs7, three demonstrated higher rosmarinic acid content than ‘Sekiho’, and two showed higher perillaldehyde content. However, none of the individuals exceeded ‘Sekiho’ for both compounds.

**Figure 4.**
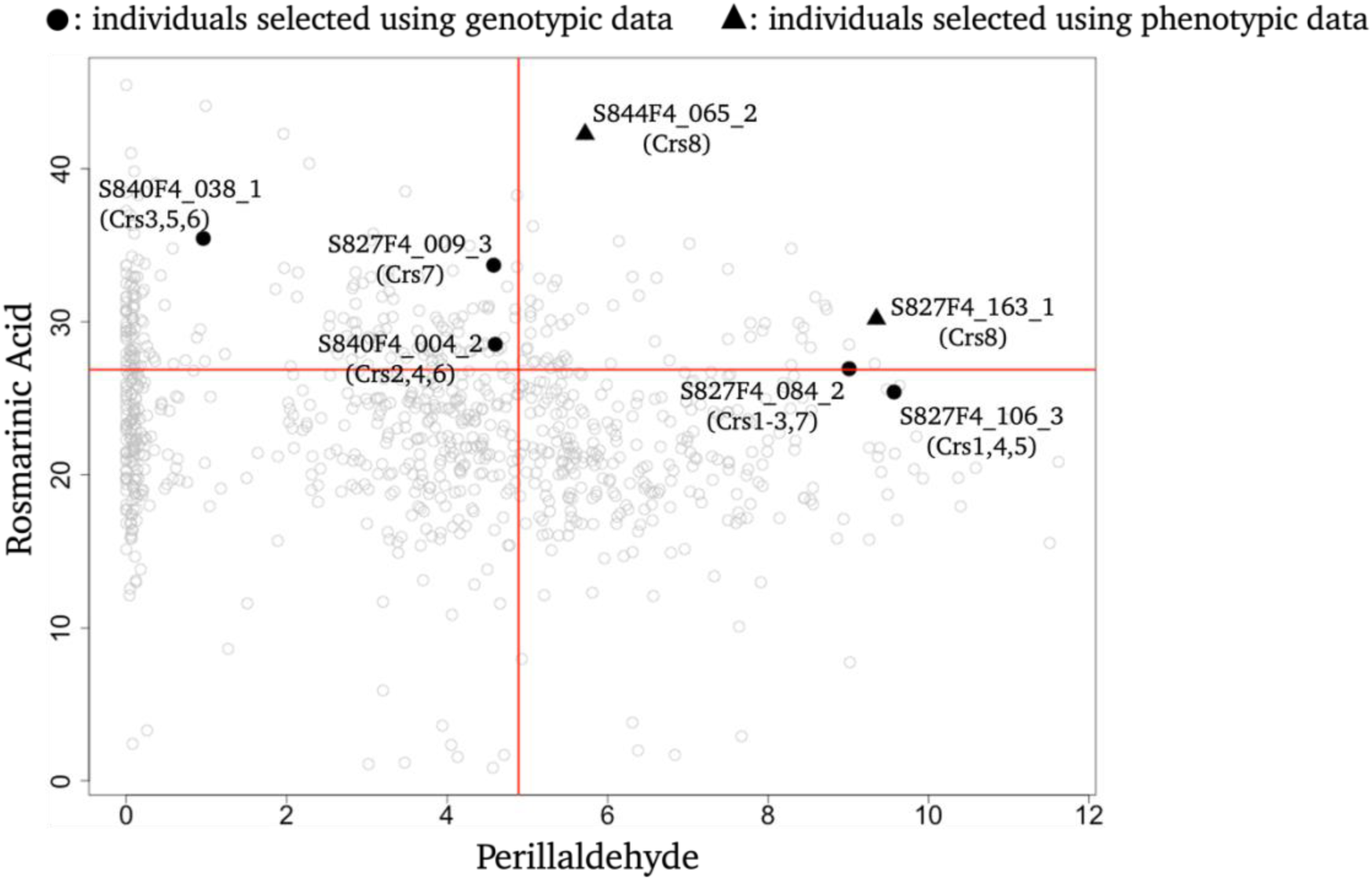
Scatter plot of perillaldehyde and rosmarinic acid phenotypes in the F₄ generation. Two red lines indicate the phenotypic values of the cultivar ‘Sekiho’ measured in the same field and in the same year. Black circles represent the five F₄ individuals used for crosses selected based on predicted additive genotypic values of G₂ progeny, whereas black triangles indicate the two individuals used for crosses selected by phenotypic selection. The seven individuals used to generate the G₂ generation are labeled with their individual IDs and the names of their corresponding crosses.

### Phenotype of the G_2_ generation

Fig. 5 shows the phenotypes of perillaldehyde, rosmarinic acid, and anthocyanins in G_2_ generation. Among the eight selected crosses, progeny derived from Crs1–Crs7 (selected based on predicted additive genotypic values of progeny) generally exhibited higher levels of perillaldehyde than those from Crs8 (selected based on phenotypes of the F_4_ generation), except for Crs4 and Crs5. Notably, Crs5 displayed a substantially lower perillaldehyde content than the other crosses across the entire G_2_ population. In addition, a bimodal distribution was observed in the progeny of Crs2–4 and Crs6, with individuals separated into two groups: those with almost no perillaldehyde, and those exhibiting values above a certain threshold. For rosmarinic acid, the mean of progeny phenotypes for Crs7 was the only one lower than that for Crs8. Considering the magnitude of variance, the progeny of Crs1–Crs7 generally showed greater variance than those of Crs8 for both perillaldehyde and rosmarinic acid, except for perillaldehyde variance in Crs5. When compared with the phenotypic values of the existing red perilla variety ‘Sekiho’, the progeny means for perillaldehyde were lower in Crs4, Crs5, Crs6, and Crs8. In contrast, the progeny means for rosmarinic acid in all eight crosses (Crs1–Crs8) exceeded that of ‘Sekiho’. For anthocyanin, the progeny means of all eight crosses were lower than that of ‘Sekiho’. The G_2_ individual with the highest sum of phenotypic values for perillaldehyde and rosmarinic acid showed 1.93-fold and 1.97-fold higher levels of these compounds, respectively, than the existing cultivar ‘Sekiho’. The mean, maximum value, and variance of the three traits in the G_2_ progeny of each cross, as well as the phenotypes of ‘Sekiho’ measured in the same field and in the same year, are summarized in Table S2.

**Figure 5.**
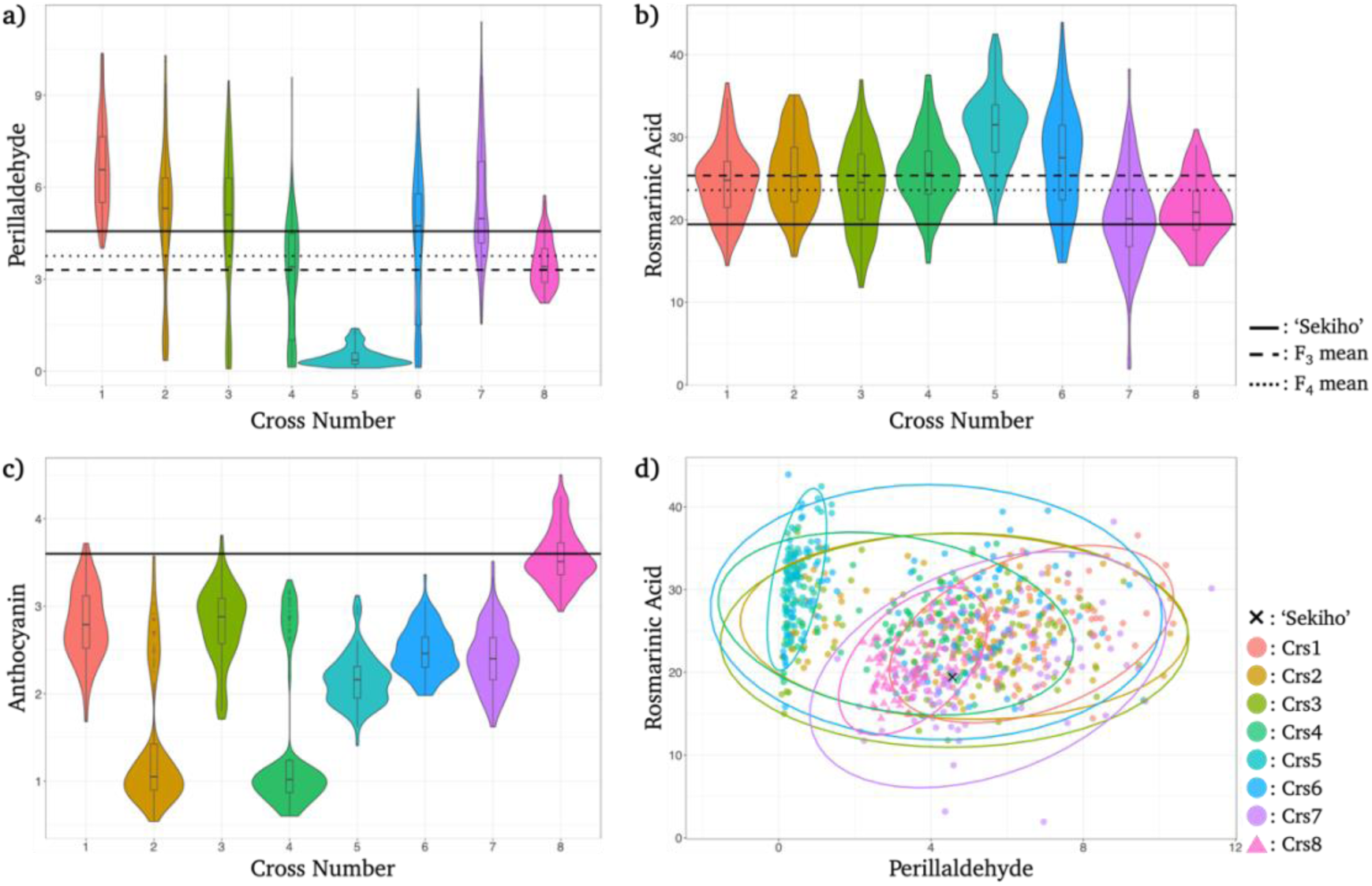
Observed phenotypes of G₂ progeny derived from each cross. Colors indicate different cross pairs. Panels (**a**)–(**c**) show violin plots of G₂ phenotypes for perillaldehyde, rosmarinic acid, and anthocyanin, respectively. In panels (**a**)–(**c**), solid lines represent the phenotypic values of the cultivar ‘Sekiho’ measured in the same field and in the same year as the G₂ generation, dashed lines indicate the mean phenotypic values of the F₃ generation, and dotted lines indicate the mean values of the F₄ generation. Panel (**d**) shows a scatter plot of perillaldehyde and rosmarinic acid phenotypes in the G₂ generation. Ellipses represent the 95% confidence intervals of the two traits for each cross pair, and crosses indicate the phenotypic values of the cultivar ‘Sekiho’ grown in the same year.

### Comparison of prediction accuracy between parental and bi-allelic models

Following the methods described in the *Inference of allele origin in the G_2_ generation* section, allele origins were inferred for the G_2_ generation. Among the 1,297 SNPs, the average number of markers for which the origin could not be determined per cross was 66.61 for Crs2, 89.31 for Crs3, 62.62 for Crs4, 61.73 for Crs5, and 57.94 for Crs8. These values represent the average of 100 progeny for each cross. For the intra-population crosses (Crs1 and Crs6–Crs7), allele origins were completely determined for all markers. Because the proportion of markers of unknown origin was small, the effects of these markers were ignored in the parental model calculations.

Fig. 6b,d show the prediction accuracies of G_2_ phenotypes using the parental model, whereas Fig. 6a,c show the results using the bi-allelic model. For both perillaldehyde and rosmarinic acid, the parental model achieved higher prediction accuracy. For perillaldehyde, the prediction accuracy increased from 0.428 to 0.513 when switching from the bi-allelic model to the parental model. For rosmarinic acid, the accuracy increased from 0.429 to 0.447, with a smaller improvement compared with perillaldehyde. The prediction accuracies for the progeny of each cross are summarized in Table S3. For perillaldehyde, the parental model demonstrated higher prediction accuracy than the bi-allelic model in all crosses except Crs5 and Crs8. In both Crs5 and Crs8, prediction accuracy was close to 0 under both models, indicating little to no predictive power. For rosmarinic acid, the parental model showed higher accuracy in Crs2–Crs4, whereas accuracy slightly decreased in the other crosses. Overall, prediction accuracy for rosmarinic acid was lower than that for perillaldehyde.

**Figure 6.**
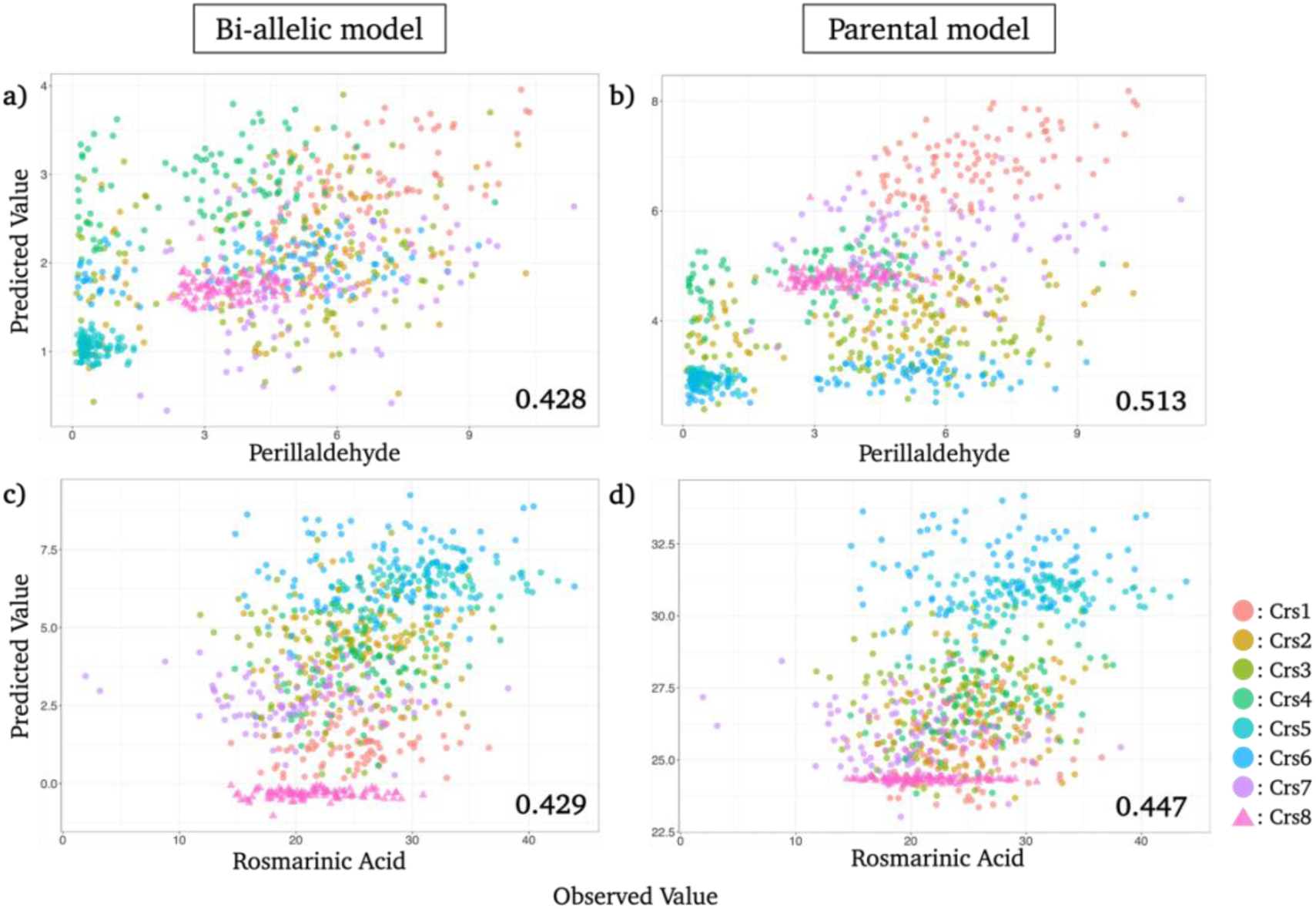
Prediction accuracy of perillaldehyde and rosmarinic acid in the G₂ generation using the bi-allelic and parental models. Colors indicate different cross pairs. The x- and y-axes represent observed and predicted values, respectively. Panels (**a**) and (**b**) show the prediction accuracy for perillaldehyde, and panels (**c**) and (**d**) show that for rosmarinic acid. Panels (**a**) and (**c**) correspond to the bi-allelic model, whereas panels (**b**) and (**d**) correspond to the parental model. The numbers in the lower right corners of each panel indicate the Pearson correlation coefficients between observed and predicted values.

## Discussion

We evaluated the effectiveness of GS in improving three major medicinal compounds in red perilla by selecting cross pairs based on the predicted additive genotypic values of the progeny and conducting actual crossing experiments. Among these compounds, perillaldehyde and rosmarinic acid were the primary targets for selection and crossing, whereas anthocyanins were targeted using a single SNP identified in a previous study. In GS-based selection, the evaluation score for each cross is defined as the predicted value of the top 1% of the progeny, enabling the method to account not only for the average additive genetic values of the two parents but also for the genetic variance generated by the cross. The results from the actual crossing experiments supported this approach; for both perillaldehyde and rosmarinic acid, most of the crosses selected based on progeny prediction (Crs1–Crs7) displayed higher average phenotypic values than the crosses selected through the phenotypic selection of two apparently superior parents (Crs8). Notably, the progeny of Crs1–Crs7 exhibited greater variance, leading to the emergence of superior individuals compared to those derived from Crs8 (Fig. 5).

A bimodal distribution of perillaldehyde was observed in the progeny of Crs2–4 and Crs6. A genome-wide association study (GWAS) conducted on the G_2_ population identified a significant peak on chromosome 7 (data not shown). This peak corresponds to one of the QTLs previously detected for perillaldehyde in the S840 population by Kinoshita et al. (2023). At this QTL, individuals homozygous for the st40 allele exhibited perillaldehyde levels close to 0. Therefore, the bimodal distribution observed in Crs2–4 (inter-population crosses between S827 and S840) and Crs6 (intra-population cross of S840) is possibly attributable to the effect of this QTL. In the case of Crs5, nearly all individuals showed very low perillaldehyde levels, most likely due to a failed cross, in which the G_2_ progeny were derived from the selfing of S840F4_038_1 rather than from a true hybrid. Cross success or failure was determined as follows: using F_4_ genotypic data, single-nucleotide polymorphisms (SNPs) fixed for different homozygous alleles between two parental individuals were identified for each cross. If heterozygous individuals were present for all SNPs in the G_2_ population, the cross was considered successful.

One possible explanation for why the progeny of Crs1–Crs7 selected by GS displayed higher mean phenotypic values than those of Crs8 selected by phenotypic selection is the advantage of incorporating genome-wide marker information into GS. GS enables the model to capture the genetic components of phenotypic variation and accumulate favorable allelic effects across the genome (Heffner et al., 2010). In contrast, phenotypic selection cannot disentangle genetic effects from environmental noise and may select individuals whose phenotypic values are upwardly biased owing to environmental errors. In addition, this advantage of GS has been demonstrated in several previous studies using real data, where greater genetic gain was achieved through GS than phenotypic selection by leveraging genomic information (Gaynor et al., 2017; Tessema et al., 2020). In the case of Crs8, one possible reason for the lack of improvement in the perillaldehyde content among its progeny was the nature of the phenotypic selection strategy used in this study. In this approach, selection was based on identifying individuals that exceeded ‘Sekiho’ in both perillaldehyde and rosmarinic acid. Although the S840 population included individuals with high rosmarinic acid content, several were excluded from phenotypic selection because of their low perillaldehyde content. However, a GWAS conducted on the G_2_ generation revealed that perillaldehyde accumulation is largely controlled by a single locus: individuals homozygous for the st40 allele exhibited nearly zero perillaldehyde (data not shown). This limitation can be overcome by crossing individuals carrying different alleles at the same locus. In GS-based selection, the effect of this locus was captured using the prediction model. Consequently, even if one parent had very low perillaldehyde content, the model could predict high perillaldehyde levels in the progeny by accounting for the complementarity between parental alleles. Crossing is a crucial step in plant breeding, and numerous methods have been proposed for selecting optimal cross pairs. The effectiveness of approaches that utilize genomic information for cross-selection has been demonstrated in both simulation and empirical studies (Zhong & Jannink, 2007; Akdemir & Sánchez, 2016; Lado et al., 2017). Our results provide empirical support for simulation-based cross-selection, demonstrating that pre-cross-prediction of progeny performance can exploit genetic variance and assemble favorable alleles across populations.

Among the three target traits, anthocyanin was the only trait for which the progeny derived from phenotypic selection outperformed those derived from GS. The Crs8 progeny exhibited both higher mean and maximum anthocyanin content than the progeny of any of the Crs1–Crs7. Although anthocyanins were not explicitly considered in the phenotypic selection process, one individual used for Crs8 belonged to the S844 population. This population had relatively low polymorphism in the F_4_ generation (579 SNPs) and possibly shared multiple alleles with the red perilla variety ‘Sekiho’. This genetic similarity may have contributed to the higher accumulation of anthocyanins, the key pigment responsible for the red–purple coloration of perilla leaves. In GS-based selection, candidate crosses are filtered using a single SNP associated with anthocyanin content, which corresponds to a major QTL distinguishing between red and green types (Zhang et al., 2021; Kinoshita et al., 2023). Although this approach successfully ensured that all G_2_ progeny exhibited the red perilla phenotype, it did not lead to further improvement in anthocyanin concentration. Given the importance of anthocyanins as pharmacologically active compounds and contributors to the appearance of Kampo formulations, further improvement of this trait is desirable. To achieve this, anthocyanins should be treated as quantitative traits, and selection strategies should shift from marker-assisted selection to GS using genome-wide information.

Because only seven cross pairs were tested, it was difficult to evaluate the accuracy of progeny-based selection using standard metrics such as correlation coefficients between predicted and observed values. Therefore, we assessed the discrepancy between the predicted and actual data by comparing the following three distributions: 1. Distribution of predicted additive genotypic values based on simulated G_2_ genotypes. These values were calculated using the simulated G_2_ genotype data from a single simulation replicate, and population-specific marker effects were estimated from the F_4_ generation. 2. Distribution of predicted additive genotypic values based on G_2_ genotype data. Allele origins were inferred from the observed G_2_ genotypes, and additive genotypic values were calculated using these inferred origins along with the marker effects estimated from the F_4_ generation. 3. The distribution of BLUPs was estimated from the observed phenotypic data of the G_2_ individuals. The results of the comparison are shown in Fig. S3. The degree of deviation among the three distributions varied depending on the trait and cross-pair. However, several general factors can be considered as causes of these discrepancies. First, the discrepancy between the distribution of the predicted additive genotypic values of the simulated and actual G_2_ progeny possibly resulted from differences in the marker genotype data, specifically from mismatches between the expected (theoretical) and actual recombination outcomes. Supporting evidence for this interpretation was obtained by comparing the LD (measured as the correlation between markers) patterns of the simulated and actual G_2_ progeny (Fig. S4). In the simulated progeny, the LD decayed smoothly with increasing map distance, whereas a high LD persisted in the actual progeny, even at longer distances. This pattern possibly reflects the effects of strong selection imposed during F_5_ generation crosses, leading to a Bulmer effect or selective sweeps (Bulmer, 1971; Kim & Nielsen, 2004). In addition, the inability to incorporate markers whose allele origins could not be determined may have contributed to the observed deviations. Second, the discrepancy between the distribution of predicted additive genotypic values based on actual G_2_ genome data and the distribution of BLUPs calculated from the phenotypic data of the G_2_ generation is possibly attributable to inaccuracies in marker effects estimated using the F_4_ generation. Although this may partly result from factors such as trait heritability and marker density, a more critical factor is that selection and crossing altered the LD relationships between SNPs and causal QTLs, rendering the marker effects estimated in the F_4_ generation less applicable to the G_2_ generation (Vitezica et al., 2011; Van Grevenhof, Van Arendonk & Bijma, 2012; Misztal et al., 2021). Because of the limited number of crosses, it was not possible to quantitatively evaluate the discrepancy between the simulated predictions and actual phenotypes. Nevertheless, based on the considerations described above, discrepancies in both mean and variance are expected to arise between the predicted and observed distributions.

In this study, breeding was initiated from three biparental populations. When intra- and inter-population crosses were performed in the F_5_ generation, the marker effects were estimated separately for each population. The founder lines of these breeding populations—‘Sekiho’ (a cultivated red perilla variety) and three genetic resources, ‘st27’, ‘st40’, and ‘st44’—belong to different botanical varieties within *Perilla frutescens*. Specifically, ‘Sekiho’, ‘st27’, and ‘st44’ are classified as *P. frutescens* (L.) Britton var. *crispa*, whereas st40 is *P. frutescens* (L.) Britton var. *frutescens*. This botanical distinction, particularly the inclusion of st40, suggests substantial differences in genetic background among the founder lines. Accordingly, differences in LD structure and allele frequencies were expected among populations. Therefore, we considered population-specific marker effects in the genomic prediction model. As shown in Fig. 6, predictions of G_2_ phenotypes using population-specific marker effects estimated from the F_4_ generation achieved higher prediction accuracies than those based on common marker effects across populations, highlighting the importance of incorporating population-specific marker effects. In addition, previous studies have reported that accounting for population- or parent-specific marker effects can increase the power of GWAS and improve genomic prediction accuracy, particularly when marker density is low (Maurer et al., 2017; Wang et al., 2022). However, population-specific marker effects can lead to unstable estimates when the number of markers in a population is insufficient. In such cases, combining populations to increase the overall dataset size has been reported to improve the prediction accuracy (De Roos, Hayes & Goddard, 2009; Technow et al., 2012). In the present study, the S844 population, which had a limited number of polymorphic markers and consequently a low prediction accuracy, was not included in the GS-based crossing and selection. These findings suggest that the incorporation of population-specific marker effects should be determined based on the characteristics and sizes of available datasets.

In the present study, we evaluated the effectiveness of GS in the breeding of red perilla, a medicinal plant, with a particular focus on selection and crossing based on predicted progeny performance. As a result, among the three major medicinal compounds, perillaldehyde and rosmarinic acid showed substantial improvement, with the best G_2_ individual exhibiting 1.93-fold and 1.97-fold higher values, respectively, than the existing cultivar ‘Sekiho’ (the best individual was defined as the one with the highest sum of phenotypic values for perillaldehyde and rosmarinic acid in the G_2_ generation). This demonstrates considerable progress toward our breeding goal of developing a cultivar superior to ‘Sekiho’. Notably, the parental lines selected based on predicted progeny performance included individuals whose phenotypic values were inferior to ‘Sekiho’ in the F_4_ generation, yet successful crosses between these lines produced superior offspring. Although a quantitative comparison between GS-based and phenotypic selection was challenging due to the limited number of cross pairs, the effectiveness of GS-based selection was clearly demonstrated in terms of the realized genetic improvement. Moreover, the practical implementation of progeny-based selection and crossing in an actual breeding context highlighted the applicability and potential of this approach. Red perilla, an underutilized crop, lacks experienced breeders, making it an ideal target for GS approaches that fully leverage genomic information. Importantly, the breeding population used in this study was relatively small in scale; yet we successfully developed individuals with markedly higher levels of key compounds than the existing variety ‘Sekiho’ using only a limited set of genetic resources. This finding underscores the considerable untapped genetic potential that may exist in underutilized crops. We believe that our results provide a model for applying GS to breeding programs for other underutilized but valuable medicinal plants that have not yet undergone systematic genetic improvement.

## Acknowledgments

Seeds of the crossed parents of the two perilla populations used in this study were provided by the GenBank Project for Agricultural Biological Resources of the National Institute of Agrobiological Sciences. We are grateful to Ms. Terue Kurosawa for cultivating the plant materials, Dr. Yoichi Aoki for measuring the medicinal compounds of perilla, and all the technical staff at TSUMURA & CO., Japan and Kazusa DNA Research Institute, Japan, for their support.

## Author contributions

All the authors contributed to the conception and design of this study. KS and HI were involved in conceptualization. Data curation was done by SK. SK and KS helped with formal analysis. TT and HI helped with funding acquisition. The investigation was done by TT. HI was involved in the project administration. TT, MS, KS, and SI contributed resources. The software was done by SK. HI helped with the supervision. SK contributed to the visualization. SK wrote the original manuscript. HI helped write, review, and edit the manuscript. All the authors have read and agreed to the published version of the manuscript.

## Data availability

Data are available from the corresponding author upon reasonable request.

