## Supplemental File for "Progeny-based genomic selection reveals untapped genetic potential in an underutilized medicinal plant, *Perilla frutescens*"

**TableS1. Individual IDs of the eight crosses used for generating the G₂ generation.**

|  | **Individual 1** | **Individual 2** |
| --- | --- | --- |
| **Crs1** | S827F5_084_2 | S827F5_106_3 |
| **Crs2** | S840F5_004_2 | S827F5_084_2 |
| **Crs3** | S840F5_038_1 | S827F5_084_2 |
| **Crs4** | S840F5_004_2 | S827F5_106_3 |
| **Crs5** | S840F5_038_1 | S827F5_106_3 |
| **Crs6** | S840F5_004_2 | S840F5_038_1 |
| **Crs7** | S827F5_009_3 | S827F5_084_2 |
| **Crs8** | S827F5_163_1 | S844F5_065_2 |

**TableS2. Phenotypic value of G_2_ generation for each cross and ‘Sekiho’.**

|  | **Perillaldehyde** | | | **Rosmarinic Acid** | | | **Anthocyanin** | | |
| --- | --- | --- | --- | --- | --- | --- | --- | --- | --- |
|  | **Mean** | **Max** | **Variance** | **Mean** | **Max** | **Variance** | **Mean** | **Max** | **Variance** |
| **Crs1** | 6.64 | 10.36 | 2.26 | 24.63 | 36.56 | 18.62 | 2.79 | 3.72 | 0.17 |
| **Crs2** | 4.87 | 10.28 | 5.60 | 25.56 | 35.13 | 20.40 | 1.39 | 3.57 | 0.57 |
| **Crs3** | 4.62 | 9.47 | 6.12 | 23.90 | 36.95 | 27.16 | 2.80 | 3.81 | 0.19 |
| **Crs4** | 3.08 | 9.58 | 3.53 | 25.88 | 37.54 | 19.83 | 1.41 | 3.30 | 0.71 |
| **Crs5** | 0.48 | 1.39 | 0.10 | 31.22 | 42.49 | 19.60 | 2.16 | 3.12 | 0.09 |
| **Crs6** | 4.14 | 9.21 | 5.70 | 27.27 | 43.91 | 38.50 | 2.49 | 3.36 | 0.07 |
| **Crs7** | 5.50 | 11.37 | 3.55 | 20.33 | 38.21 | 33.02 | 2.42 | 3.51 | 0.13 |
| **Crs8** | 3.53 | 5.72 | 0.62 | 21.38 | 30.92 | 12.97 | 3.57 | 4.50 | 0.10 |
| **‘Sekiho’** | 4.56 | | | 19.43 | | | 3.60 | | |

**TableS3. Prediction accuracy for each cross using bi-allelic or parental model in each trait.**

|  | **Perillaldehyde** | | **Rosmarinic Acid** | |
| --- | --- | --- | --- | --- |
|  | **Bi-allelic model** | **Parental model** | **Bi-allelic model** | **Parental model** |
| **Crs1** | 0.501 | 0.533 | 0.062 | 0.039 |
| **Crs2** | 0.295 | 0.342 | 0.084 | 0.216 |
| **Crs3** | 0.187 | 0.264 | 0.088 | 0.154 |
| **Crs4** | 0.237 | 0.337 | 0.281 | 0.355 |
| **Crs5** | -0.067 | -0.093 | 0.14 | 0.108 |
| **Crs6** | 0.361 | 0.48 | 0.196 | 0.165 |
| **Crs7** | 0.208 | 0.218 | -0.012 | -0.041 |
| **Crs8** | 0.054 | 0.003 | 0.085 | -0.028 |

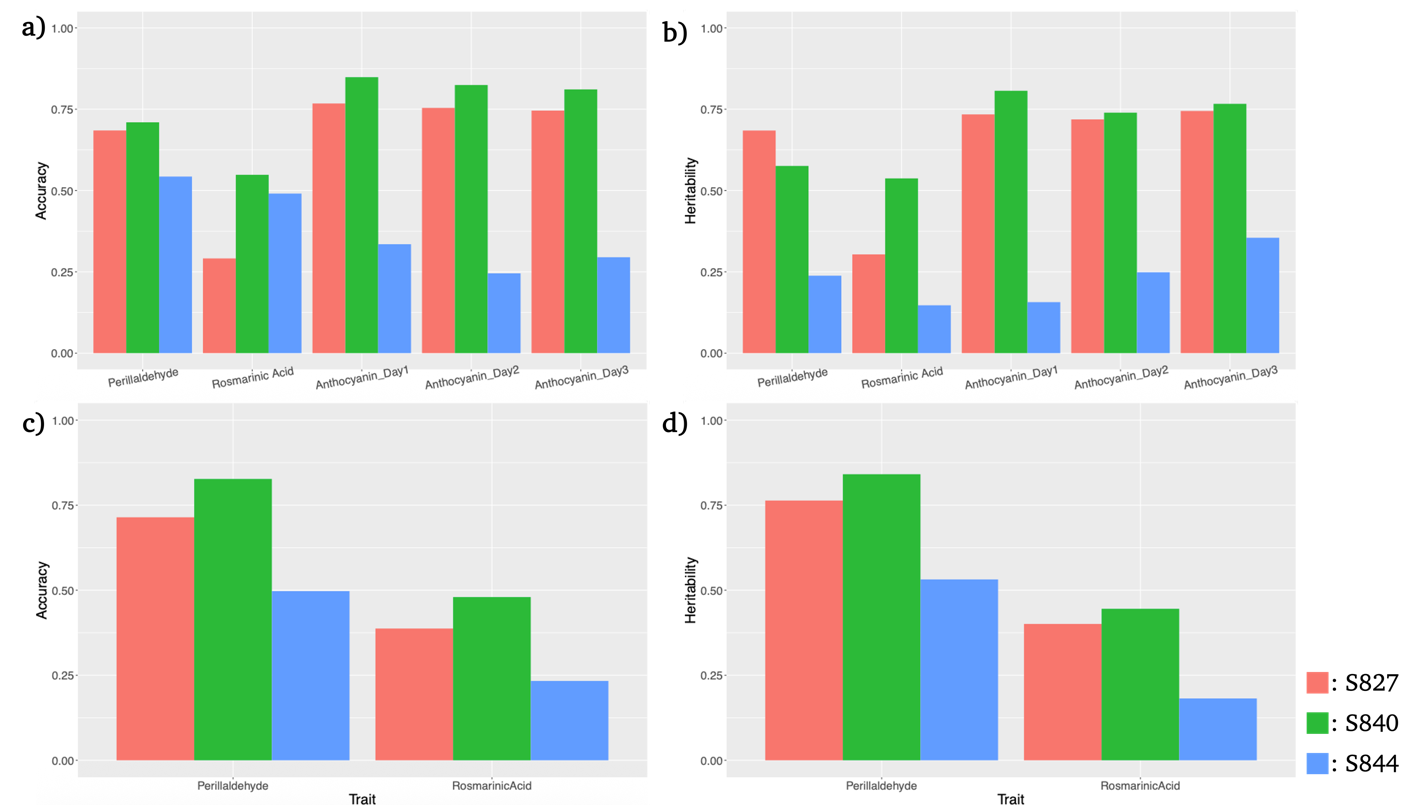

**FigureS1.** Genomic prediction (GP) accuracy and genomic heritability for each trait in the F₃ and F₄ generations estimated using the BayesB model with ‘Sekiho’ as the reference genome. Panels (a) and (b) show the results for the F₃ generation, and panels (c) and (d) show those for the F₄ generation. Panels (a) and (c) indicate GP accuracy, while panels (b) and (d) indicate genomic heritability. Red, green, and blue bars represent the S827, S840, and S844 populations, respectively. Traits measured in the F₃ generation were perillaldehyde, rosmarinic acid, and anthocyanin (three measurement dates), whereas in the F₄ generation, perillaldehyde and rosmarinic acid were measured.

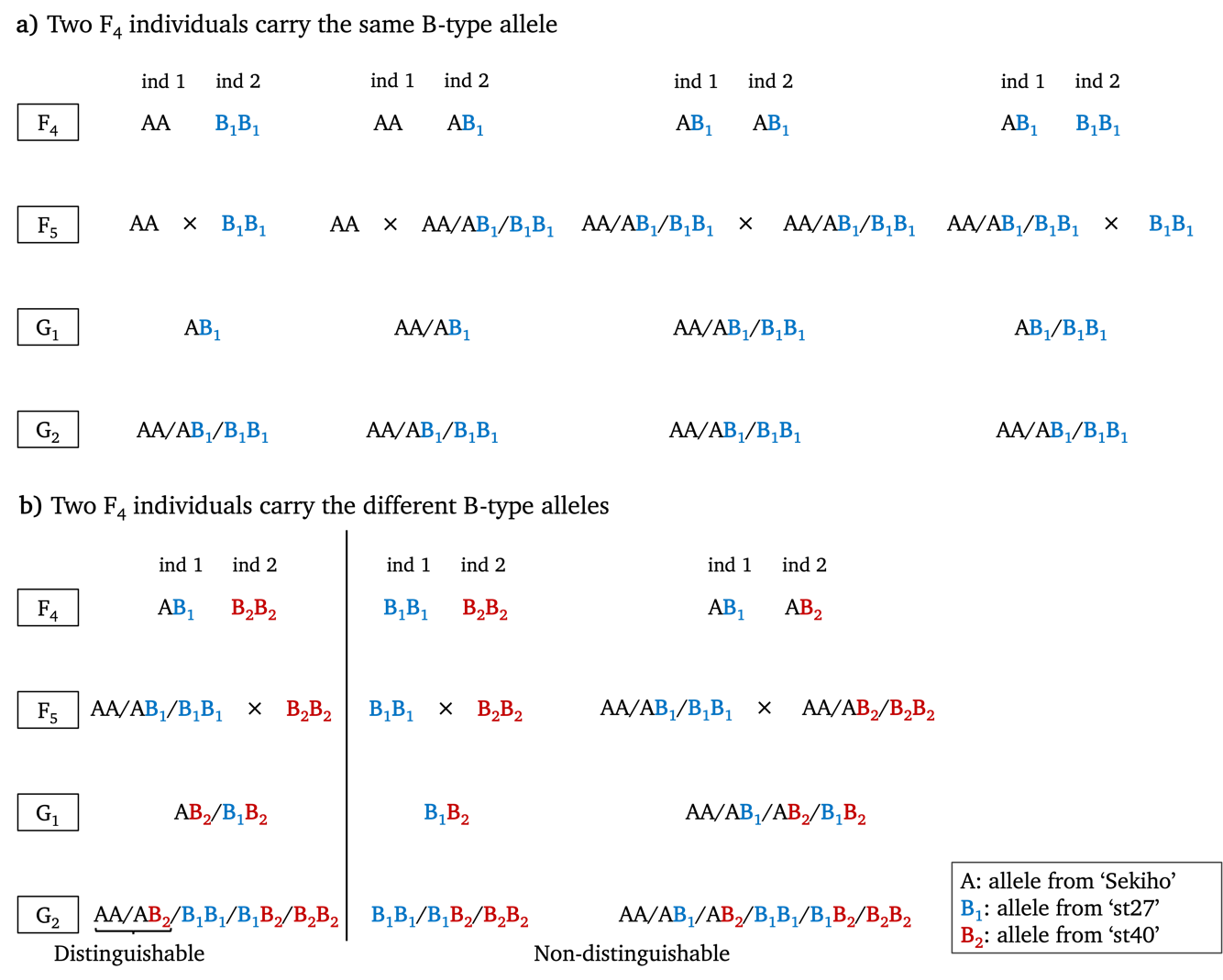

**FigureS2.** Schematic representation of the method used to infer allele origin in the G₂ generation. A denotes alleles derived from ‘Sekiho’, B₁ from ‘st27’, and B₂ from ‘st40’. Panel (a) illustrates cases where two F₄ individuals possessed B alleles of the same origin, whereas panel (b) shows cases where the two individuals carried B alleles of different origins. For each SNP genotyped in the F₄ generation, the combinations of genotypes between the two individuals constituting a cross are shown. The combination AA–AA is omitted because it necessarily results in an AA genotype in the G₂ generation. Patterns symmetric with respect to B₁ and B₂ are also not shown.

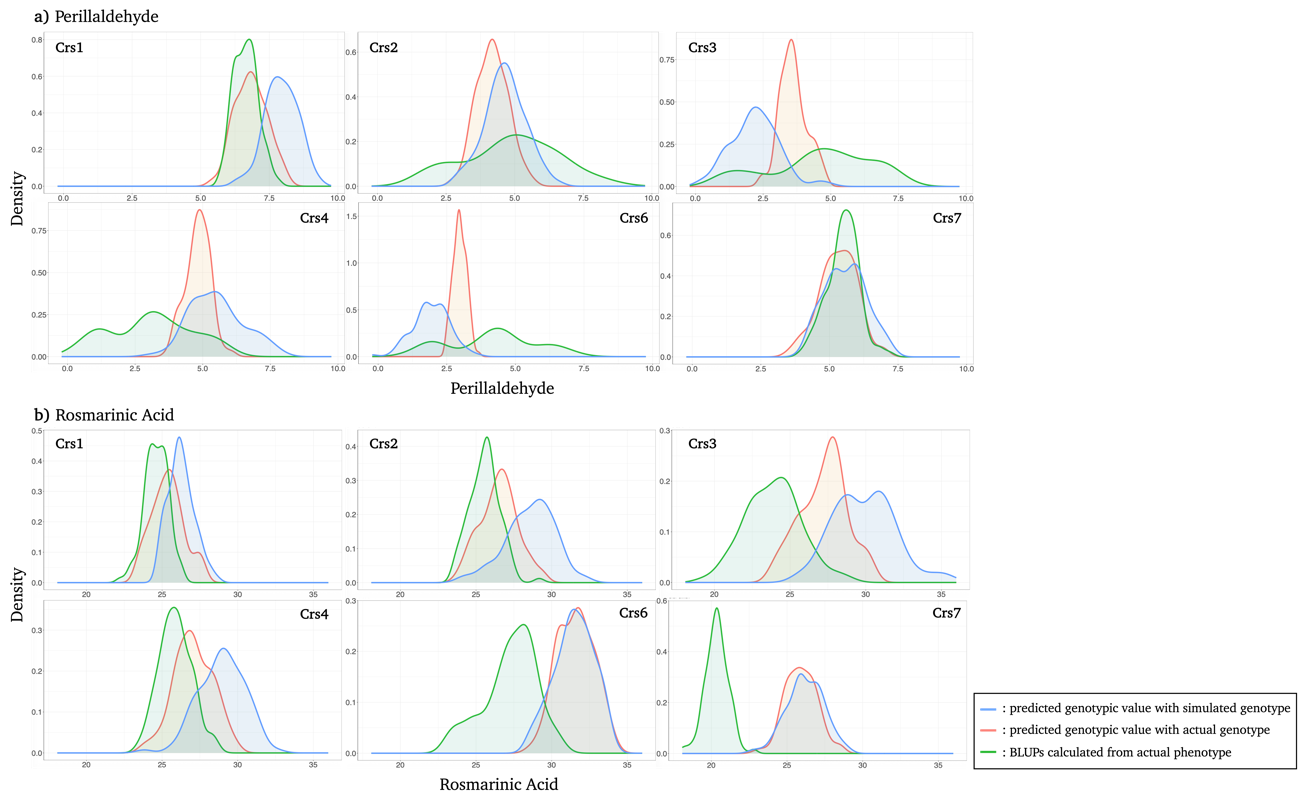

**FigureS3.** Distributions of predicted additive genotypic values and BLUPs of the G₂ generation for each cross. Crs5 was removed due to the failure of crossing. Panel (a) shows the results for perillaldehyde, and panel (b) for rosmarinic acid. Blue distributions represent predicted additive genotypic values based on simulated genotype data, red distributions represent predicted additive genotypic values based on actual G₂ genotype data with inferred allele origins, and green distributions represent BLUPs of the G₂ generation calculated using actual G_2_ phenotype data.

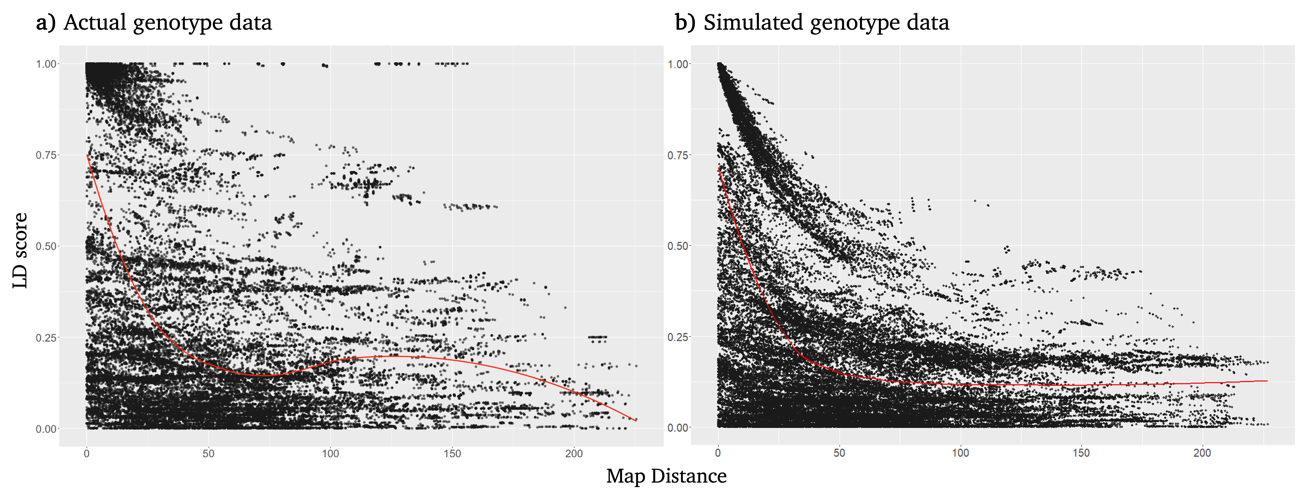

**FigureS4.** Linkage disequilibrium (LD) in the G₂ generation based on actual and simulated genotypic data. Panel (a) shows LD calculated from the actual genotypic data, and panel (b) shows LD calculated from the simulated genotypic data. LD was estimated as the correlation between pairs of markers. The red line represents the smoothed trend.
